# Functional characterization of the 26RFa/orexin neurons of the lateral hypothalamic area reveals their key role in the control of glucose homeostasis

**DOI:** 10.64898/2026.10.13.758728

**Authors:** Mélodie Devère, Caroline Lusurier, Saloua Takhlidjt, Yamina Cherifi, Emmanuelle Nédélec, Alexandre Bénani, Marius Rosay, David Godefroy, Arnaud Arabo, Benjamin Lefranc, Jérôme Leprince, Gaëtan Prévost, Nicolas Chartrel, Marie Picot

**Affiliations:** Univ Rouen Normandie, Inserm, Normandie Univ, NorDiC UMR 1239, F-76000 Rouen, France; Centre des Sciences du Goût et de l’Alimentation (CSGA), Université Bourgogne Europe, Institut Agro Dijon, CNRS, INRAE, 21000 Dijon, France; Univ Rouen Normandie, Normandie Univ, INSERM US 51, CNRS UAR 2026, HeRacLeS, F-76000 Rouen, France; Univ Rouen Normandie, Inserm, Normandie Univ, NorDiC UMR 1239, CHU Rouen, Department of Endocrinology, Diabetes and Metabolic Diseases, F-76000 Rouen, France

**Keywords:** neuroendocrinology, glucose homeostasis, energy metabolism, hypothalamus, diabetes, neuropeptide

## Abstract

Defining the nature and molecular mechanisms of the neurocircuitry controlling glucose homeostasis is crucial in understanding how alterations in these pathways promote the development of obesity and type 2 diabetes. Here, we functionally characterize a population of lateral hypothalamic neurons co-expressing the anti-hyperglycemic neuropeptides 26RFa and orexin. Using complementary chemogenetic, pharmacological and genetic approaches, including mice deficient for 26RFa, orexin or both neuropeptides, we investigated the respective contribution of each signaling system to glucose homeostasis.

We found that lateral hypothalamic 26RFa neurons extensively co-express orexin, defining a distinct peptidergic population with convergent anti-hyperglycemic properties. Unexpectedly, chemogenetic activation of these neurons potentiated glucose-induced hyperglycemia despite the anti-hyperglycemic properties of both neuropeptides. To understand this paradox, we examined the transcriptional responses of the 26RFa and orexin peptidergic systems and found that neuronal activation was associated with coordinated, glycemic state-dependent downregulation of both signaling systems. Furthermore, genetic loss-of-function approaches revealed distinct contributions of the two neuropeptidergic systems, with orexin signaling mediating most of the metabolic consequences of neuronal activation, whereas 26RFa signaling is required for the dynamic regulation of orexin expression.

Finally, our findings provide new insights into the functional organization of lateral hypothalamic 26RFa/orexin neurons and advance our understanding of how coordinated neuropeptidergic signaling contributes to systemic glucose homeostasis.

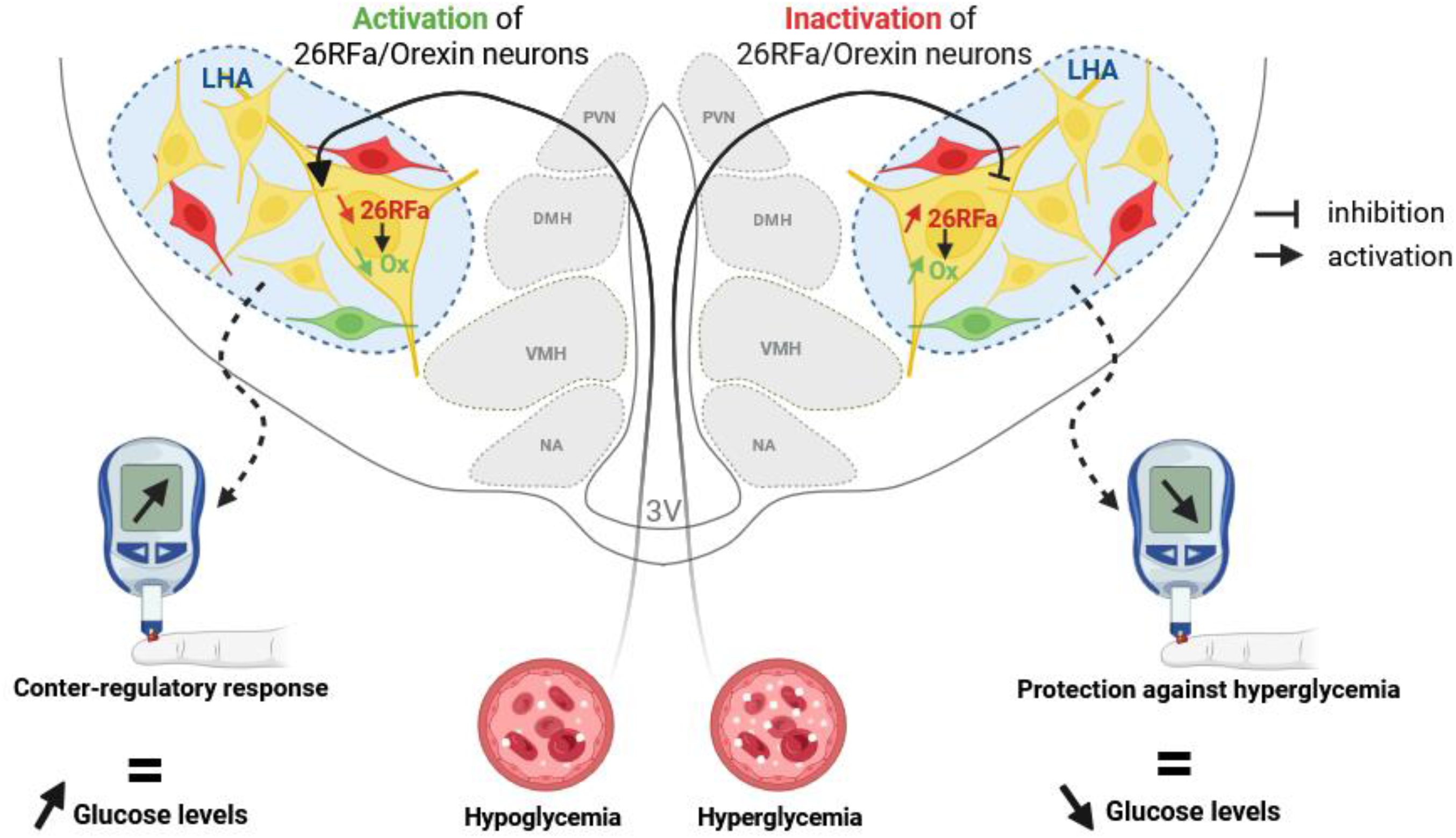

## 1. Introduction

The hypothalamus is a critical component of the homeostatic system notably by regulating glucose metabolism. Indeed, it is now widely accepted that the control of glucose homeostasis results from a complex and highly coordinated interaction between the brain and the pancreatic islets as it is suspected the brain accounts for 50% of overall glucose disposal and can normalize glycemia in rodent models of diabetes (Carey *et al*, 2013; Fujikawa *et al*, 2013). Defining the nature and molecular mechanisms of the neurocircuitry controlling glucose homeostasis is crucial in understanding how alterations in these pathways promote the development of major associated pathologies such as type 2 diabetes and would nurture the development of alternative therapeutic strategies to treat this disease that has become an epidemic worldwide issue (Brüning & Fenselau, 2023). Recent development of sophisticated approaches such as chemogenetics and genome-wide association study has enable the identification of newly neuronal subpopulations and their functional role in the control of glucose metabolism but also their activity in freely behaving animals (Brüning & Fenselau, 2023; Haspula & Cui, 2023). As a contribution to this common effort, the present study focuses on a subpopulation of neurons located in the lateral hypothalamic area (LHA), that co-express the neuropeptides 26RFa and orexin, because a number of evidence suggest that the 26RFa/orexin neurons of the LHA may be important players of the central regulation of glucose homeostasis.

26RFa (also referred to as QRFP) is a hypothalamic neuropeptide discovered concurrently by us and others (Chartrel *et al*, 2003; Fukusumi *et al*, 2003; Jiang *et al*, 2003), and identified as the cognate ligand of the human orphan G protein-coupled receptor GPR103 (Chartrel *et al*, 2003; Fukusumi *et al*, 2003; Jiang *et al*, 2003; Leprince *et al*, 2017; Takayasu *et al*, 2006) that regulates energy and glucose metabolism (Devère *et al*, 2024). Indeed, i.c.v. administration of 26RFa robustly stimulates food intake (Chartrel *et al*, 2003; Lectez *et al*, 2009; Moriya *et al*, 2006; Takayasu *et al*, 2006), and this orexigenic activity is mediated by the NPY and POMC neurons in the arcuate nucleus (Arc) (Lectez *et al*, 2009). Moreover, we recently reported that i.c.v injection of 26RFa, induces a sustained anti-hyperglycemic effect associated with an increase of insulin production by the pancreatic islets (El Mehdi *et al*, 2022). Finally, we have provided conclusive evidence that the 26RFa/GPR103 peptidergic system is a key relay for the central regulation of glucose metabolism by insulin (El Mehdi *et al*, 2022).

The orexin system comprises the two neuropeptides orexin A and orexin B (also designed hypocretin-1 and hypocretin-2), encoded by the same polypeptide precursor, and their two receptors orexin/hypocretin 1 (OX1R) and orexin/hypocretin 2 (OX2R) (de Lecea *et al*, 1998; Sakurai *et al*, 1998). Interestingly, orexins were initially recognized as regulator of feeding behaviour in the brain by stimulating appetite (Sakurai *et al*, 1998). Subsequently, the observation that narcolepsy in human and animals is caused by deficiency of the orexin system highlighted the key role played by orexins in arousal and sleep/wake states (Chemelli *et al*, 1999; Lin *et al*, 1999; Seifinejad *et al*, 2023; Thannickal *et al*, 2000). In addition, emerging evidence suggest that orexin-producing neurons are involved in the control of glucose homeostasis by modulating glucose production and utilization in the peripheral tissues notably via the autonomic nervous system (Tsuneki *et al*, 2012). It was also found that orexin-neurons of the LHA are glucose-inhibited and thus that low glucose concentrations or hypoglycemia activate this neuronal population (Burdakov & González, 2009; Paranjape *et al*, 2006; Sheng *et al*, 2014). Moreover, we recently showed that orexin deficiency induces a decrease of glucose tolerance and insulin sensitivity (Devère *et al*, 2025).

The recent identification of lateral hypothalamic neurons co-expressing 26RFa and orexin raises fundamental questions regarding the functional significance of this neuropeptidergic population. Because both neuropeptides have been implicated in the central regulation of energy and glucose homeostasis, their coexistence within the same neurons suggests a previously unexplored mode of neuropeptidergic integration within the lateral hypothalamus. Whether these two peptidergic systems operate independently, redundantly or through hierarchical interactions within the same neuronal population, and how each contributes to glucose homeostasis, remains completely unknown. Addressing these questions is essential to understand how multiple neuropeptidergic signaling are integrated within hypothalamic circuits controlling glucose homeostasis. Here, we combined complementary neuroanatomical, chemogenetic, pharmacological and genetic approaches, including 26RFa-, orexin- and double-knockout mouse models, to characterize this lateral hypothalamic 26RFa/orexin neuronal population, determine the functional organization of its co-expressed peptidergic systems and define their respective contributions to the central regulation of glucose homeostasis.

## 2. Methods

### 2.1. Animals

26RFa^+/+^, 26RFa^+/−^, 26RFa^−/−^, Orexin^−/−^, 26RFa^+/−^ Orexin^−/−^, 26RFa^−/−^ Orexin^−/−^ and 26RFa^+/−^ -tdTomato male C57Bl/6J mice (2–4 months old), weighing 22–35 g, were housed with free access to standard diet (UAR, Villemoisson-sur-Orge, France) and tap water. 26RFa^−/−^ (iCre knock-in) mice were obtained from Sakurai (International Institute for Integrative Sleep Medecine, Tsukuba, Ibaraki, Japan) as previously described (El Mehdi *et al*, 2020). tdTomato mice, which express robust tdTomato fluorescence following Cre-mediated recombination, were purchased from The Jackson Laboratory (B6.Cg-*Gt(ROSA)26Sor^t14(CG6-tdTomato)Hze^*, JAX stock #007914, Bar Harbor, ME), and have been previously described (Madisen *et al*, 2010). 26RFa^+/-^-tdTomato mice were generated by crossing 26RFa^+/−^ (iCre knock-in) mice with tdTomato mice. Orexin^−/−^ mice were obtained from The Jackson laboratory (Strain #:008867, RRID:IMSR_JAX:008867) as previously described (Devère *et al*, 2025). 26RFa^+/−^ Orexin^−/−^, 26RFa^−/−^ Orexin^−/−^ were generated by crossing 26RFa^+/−^ (iCre knock-in) mice with Orexin^+/−^ mice. All mice were kept in a ventilated room at a temperature of 22±1°C under a 12-hour light/12-hour dark cycle (light on between 7:00 and 19:00). All experiments were carried out between 07:30 and 18:00 in the testing rooms adjacent to the animal rooms. Mice were housed at three to five per cage. Unless otherwise stated, all tests were conducted with naïve cohorts of mice that were habituated to physiological and behavior protocols before the beginning of experiments.

All experimental procedures were approved by the Normandy Regional Ethics Committee (authorisation: APAFIS no. 2022090217265485) and were carried out in accordance with Council Directive 86/609/EEC of 24 November 1986.

### 2.2. RNAscope in situ hybridization procedure

26RFa^+/+^ and 26RFa^-/-^male mice were deeply anaesthetized and perfused transcardially with sterile PBS followed by sterile ice-cold phosphate-buffered 4% paraformaldehyde. The brains were removed, post-fixed overnight at 4°C in sterile 4% paraformaldehyde and transferred to sterile 15% sucrose (12 h) and then to sterile 30% sucrose (12 h) in 0.1 mol/L PBS (pH 7.4) at 4°C. Finally, 12-μm-thick sections, collected every 40 μm over the whole length of the hypothalamus (15 sections/hypothalamus), were cut on a cryostat (CryoStar™ NX70) and mounted on SuperFrost Plus slides (Thermo Fisher Scientific, Illkirch, France) and subsequently stored at −80°C. Fluorescent *in situ* hybridization for the simultaneous detection of 26RFa and Orexin transcripts was performed using an RNAscope assay (Advanced Cell Diagnostics [ACD], Rennes, France). The 26RFa probe targeted the region 112–1081 (accession number NM_183424.4; ACD, Cat no. 464341-C3). The Orexin probe targeted the region 2-577 (accession number NM_010410.2; ACD, Cat no. 490461-C2). The GPR103 probe targeted the region 2-1148 (accession number NM_198192.2; ACD, Car no. 566341). Negative and positive control probes recognizing dihydrodipicolinate reductase and DapB (a bacterial transcript) and PolR2A (C1 channel), PPIB (C2 channel) and UBC (C3 channel), respectively, were processed in parallel with the target probes to ensure tissue RNA integrity and optimal assay performance. In addition, the specificity of the 26RFa probe was previously described in brain sections of 26RFa^−/−^ mice (El Mehdi *et al*, 2022). All incubation steps were performed at 40°C using a humidified chamber and a HybEz oven (ACD). On the day of the assay, all the hybridization steps (i.e. target retrieval, dehydration, probe hybridization, amplification steps and detection of the probe) were performed according to the online protocol for the RNAscope multifluorescent assay. TSA Vivid Fluorophore 520 (PN 323271, ACD), TSA Vivid Fluorophore 650 (PN 323273, ACD) and TSA Vivid Fluorophore 570 (PN 323271, ACD) were respectively used to detect orexin, 26RFa and GPR103 mRNA.

### 2.3. Immunohistochemistry procedure

26RFa^+/-^-tdTomato male mice were deeply anaesthetized and perfused transcardially with PBS followed by ice-cold PBS 4% paraformaldehyde. The brains were removed, post-fixed overnight at 4°C in sterile 4% paraformaldehyde and transferred into PBS. Brain sections (30 µm thick) were performed with a vibratome (VT1000 S 2018, Leica) before being submitted to immunohistochemistry. HCRT antibody 1:500 (mouse monoclonal HCRT, Santa Cruz Biotechnology, SC-80263, Dallas, TX, USA) was applied during 48h at 4°C. After rinsing with 1xPBS (15 min), Alexa-conjugated antibodies (donkey anti-mouse-488 (1:300), Invitrogen Life Technologies, CA, USA) was applied during 2h at room temperature. NeuN antibody 1:500 (rabbit polyclonal NeuN, Abcam, Ab104225, Cambridge, UK) was applied 48h at 4°C. After rinsing with 1xPBS (15 min), Alexa-conjugated antibodies (donkey anti-rabbit-647 (1:500), Invitrogen Life Technologies) was applied during 2h at room temperature. C-fos antibody 1:1000 (rabbit polyclonal C-fos, Abcam, Ab190289) was applied during 48h at 4°C. After rinsing with 1xPBS (15 min), Alexa-conjugated antibodies (donkey anti-rabbit-647 (1:300), Invitrogen Life Technologies) was applied during 2 h at room temperature.

### 2.4. Microscopy

Images were captured using a Leica Thunder Imager 3D (DM6 B; Leica, Paris, France) with a 20× and 100× objective. Maximum intensity projections were made in Fiji/Image J V1.53q (National Institutes of Health, Washington, DC, USA) and images were adjusted for brightness and contrast. Quantitative analysis of the 26RFa-expressing neurons number in hypothalamic sections was performed also using Fiji/Image J V1.53q.

### 2.5. Stereotaxic viral injection

26RFa^+/−^, 26RFa^−/−^, 26RFa^+/−^ Orexin^−/−^ and 26RFa^−/−^ Orexin^−/−^ male mice (8-10 weeks) were anaesthetized with isoflurane/buprenorphin and placed in a stereotaxic frame. Cre-dependent adeno associated virus injected were pAAV-hSyn-DIO-hM3D(Gq)-mCherry (Addgene plasmide #44361), pAAV-hSyn-DIO-hM4D(Gi)-mCherry, (Addgene Plasmid #44362), for chemogenetic activation and inhibition, respectively, and pAAV-hSyn-DIO-mCherry (Addgene Plasmid #50459). 200nL of AAV (at the rate of 40nL/min) were bilaterally injected using a 5µL Hamilton neuros syringe (33gauge, outer diameter 0.21 mm; REF#65460-02 Hamilton, Reno, NV, USA) into the LHA (AP – 0.85 mm, ML ± 0.90 mm, − 5.60 mm from the surface of the skull). Mice were allowed 2 weeks for recovery and viral expression before to be habituated to handling by investigators. DREADD expression in the LHA was verified at the end of the experiments by the expression of mCherry fluorescent tag included in the DREADD vector. The images presented in suppl Fig. 1 clearly show that the distribution of the DREADD vector is very similar to the distribution of 26RFa mRNA using the RNAscope approach and the distribution of tdTomato in 26RFa-tdTomato mice that express the fluorescent protein specifically in 26RFa-expressing cells (Picot *et al*, 2025) (suppl Fig. 1 B-D).

### 2.6. Central administration of drugs

Test substances, including 26RFa (5.10^-4^M, produced by the HeRacLeS platform at the University of Rouen, France), the GPR103 antagonist 25e (10^−4^ M, Custom synthesized by AstraZeneca), Orexin-A (10^-6^M, orb372620, Clinisciences, France), and OxR antagonist suvorexant (10^−4^ M, MK-4305, Clinisciences, France), were dissolved in HEPES buffer and injected intracerebroventricularly (i.c.v.) in a volume of 2 μL (1µL/min). To perform the i.c.v. administration, mice were anaesthetized with isoflurane and placed in a stereotaxic frame. A stainless steel 26 gauge guide cannula (Phymep, Paris, France) was inserted into the right lateral ventricle (0.8 mm lateral to the bregma and 2.2 mm ventral to the dura mater). A stainless steel dummy small cap for 26 gauge cannulae (Phymep, Paris, France) was inserted to prevent occlusion of the guide cannula and leakage of cerebrospinal fluid. The cannula placement was secured with dental acrylic cement. Mice were allowed 1 week for recovery before to be habituated to handling by investigators. The i.c.v. injections were performed using a Hamilton syringe (outer diameter 0.5 mm; Hamilton, Reno, NV, USA). The injections were realized under anaesthesia, which was induced by i.p. administration of diazepam (5 mg/kg) and ketamine (100 mg/kg).

### 2.7. Blood glucose and insulin measurements

For measurements of basal glycemia, glucose tolerance, pyruvate tolerance and insulinemia, mice were fasted for 16 h before the test. ip administration of clozapine-N-oxide (CNO; Sigma Aldrich, Saint Louis, MO, USA) at a dose of 0.1 mg/kg or vehicle was performed 1h before the beginning of the different glycemic tests, and each test was performed twice to ensure that each mouse serves as its own chemogenetic modulation control. Specific activation of the 26RFa-expressing neurons of the LHA after injection of CNO was verified by visualization of cFos expression in the LHA showing that the number of AAV- expressing neurons co-expressing cFos is significantly increased in the LHA of mice injected with CNO compared with the mice injected with hepes (suppl Fig. 1E). For glucose tolerance test, mice were injected (i.p.) with glucose (2 g/kg). For pyruvate tolerance test, mice were injected (i.p.) with sodium pyruvate (2 g/kg; Sigma Aldrich, Saint Louis, MO, USA). For insulin tolerance test, mice were fasted for 6 h before the test with free access to water, and then injected (i.p.) with 0.75 units/kg of human insulin (Eli Lilly, Neuilly-sur-Seine, France). Plasma glucose concentrations were measured from tail vein samplings at various times using an Accu-Chek Performa glucometer (Roche Diagnostics, Saint-Egreve, France). Plasma insulin concentrations were determined using an ultrasensitive insulin AlphaLisa Detection Kit (AlphaLISA Human and Mouse Insulin Kit, cat number AL3184C, Revvity, Waltham, MA, USA).

### 2.8. Quantitative PCR

Total RNA from hypothalami was isolated with Nucleospin RNA Kit (MACHEREY-NAGEL). Relative expression of 26RFa, GPR103, orexin, OX1R and OX2R genes was quantified by real-time PCR with appropriate primers (Table 1). β-actin and GADPH were used as internal control for normalization. PCR was carried out using Gene Expression Master Mix 2X assay (Applied Biosystems, Courtaboeuf, France) in an ABI Prism 7900HT Fast Real-Time PCR System (Applied Biosystems). The purity of the PCR products was assessed by dissociation curves. The amount of target cDNA was calculated by the comparative threshold (Ct) method and expressed by means of the 2-ΔΔCt method.

**Table 1.**
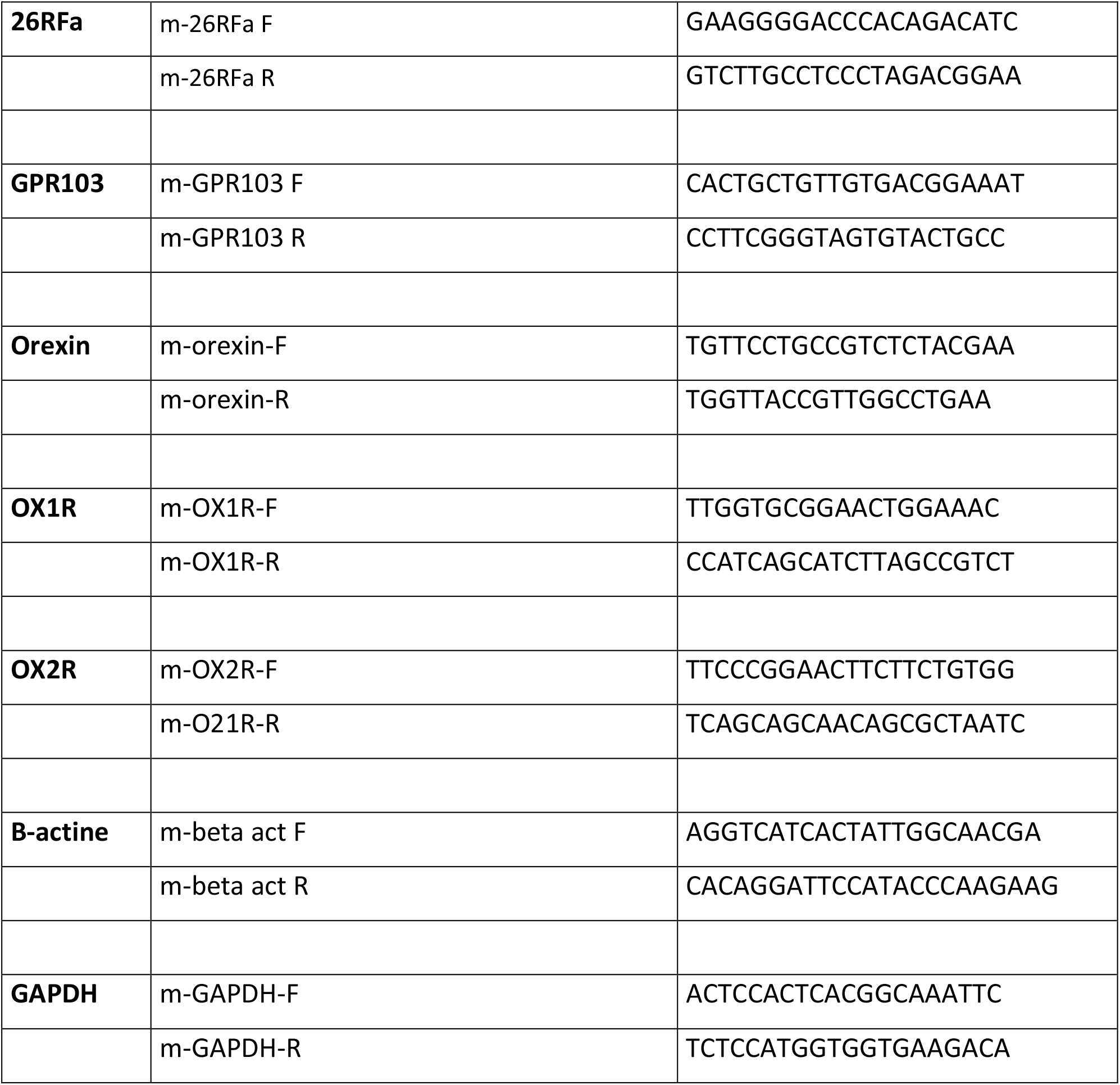

### 2.9. Statistical analysis

Statistical analysis was performed with GraphPad Prism (V.10). A Student’s t-test, a one-way analysis of variance (ANOVA) was used for comparison between the groups. A two-way ANOVA was used for quantitative PCR analysis. A post-hoc comparison using Tukey HSD was applied according to the one and two-way ANOVA results. All p values ≤0.05 were considered significant (*p ≤ 0.05, **p ≤ 0.01, ***p ≤ 0.001 and ****p ≤ 0.0001).

## 3. Results

### 3.1. Definition of a 26RFa/orexin neuronal population in the lateral hypothalamus

Distribution of 26RFa mRNA in the hypothalamus (from Bregma −1.06 to −1.70 mm) using RNAscope *in situ* hybridization technology indicated the presence of a large population of 26RFa-expressing cells in the LHA and a smaller population of positive cells located ventrally (Fig. 1A). Double-labeling experiments showed an important co-expression of 26RFa and orexin mRNAs in the same cells of the LHA (Fig. 1A, B). Indeed, quantitative analysis revealed that 90% of the 26RFa-positive cells express orexin mRNA and, conversely, 86% of the orexin-expressing cells co-express 26RFa mRNA (Fig. 1B). This molecular overlap was confirmed by immunohistochemical analysis of hypothalamic slices from 26RFa-tdTomato mice, showing important co-localization between tdTomato and orexin A antibodies (Fig. 1C, D). Quantitative analysis revealed that 86% of tdTomato cells are labelled by orexin-A antibodies and, conversely, 62% of the orexin-immunostained cells correspond to tdTomato cells (Fig. 1D). Finally, NeuN labelling further indicated that 66% of 26RFa/orexin expressing cells present in the LHA correspond to neurons (Fig. 1E). Together, these findings identify a previously uncharacterized lateral hypothalamic neuronal population co-expressing 26RFa and orexin, raising the question of whether these two neuropeptides exert complementary or distinct functions in the regulation of glucose homeostasis.

**Figure 1.**
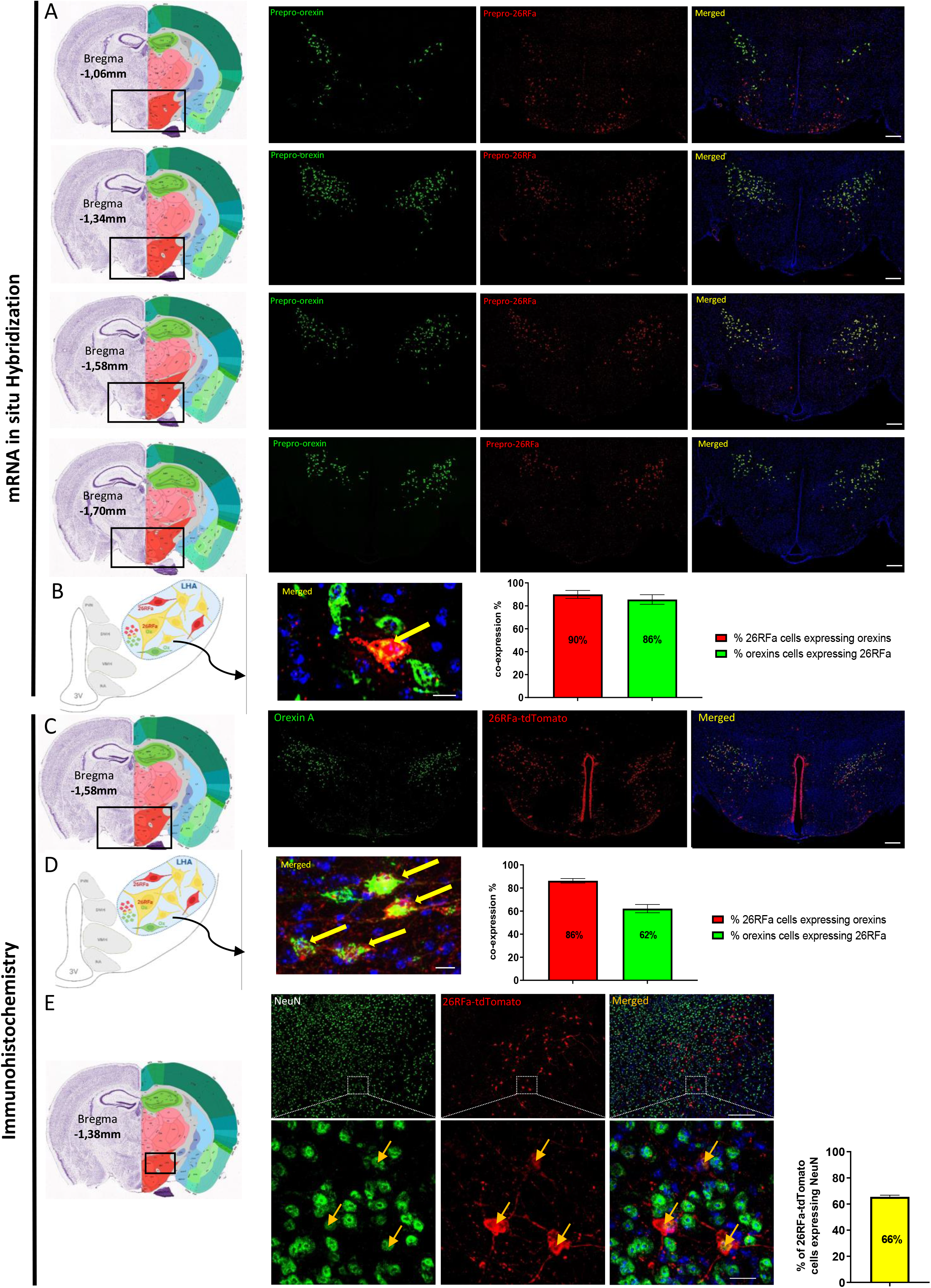
Lateral hypothalamic 26RFa-expressing neurons predominantly co-express orexin. **A)** Panoramic views of hypothalamic antero-posterior sections (Bregma −1.06 to −1.70 mm, schematic coronal sections adapted from the Allen Mouse Brain Atlas) showing the distribution of *prepro-orexin* (green) and *prepro-26RFa* (red) mRNAs in the LHA, as revealed by RNAscope in situ hybridization. **B)** Representative photomicrograph of a cell in the LHA co-expressing *prepro-orexin* and *prepro-26RFa* (yellow arrow), along with the quantitative analysis of cells co-expressing 26RFa and orexin in the LHA. Data are presented as mean ± SEM from 5 wild-type male mouse hypothalami, with 12 to 15 sections analyzed per animal. **C)** Panoramic view of a hypothalamic section of a 26RFa-tdTomato (red) mouse brain immunostained with orexin A antibody showing the extensive co-labelling of the orexin-A antibody with tdTomato. **D)** Higher magnification image showing 26RFa-tdTomato cells immunolabeled with orexin A in the LHA (yellow arrow), along with the quantitative analysis of double-labeled cells. Data represent mean ± SEM from 4 wild-type male mouse hypothalami, with 12 to 15 sections per animal. **E)** Panoramic and high-magnification views of LHA sections from 26RFa-tdTomato mice (red) immunostained for the neuronal marker NeuN (green) showing the co-localization of the NeuN-positive and tdTomato-positive cells (yellow arrow), along with the quantitative analysis of double-labeled cells. Data represent mean ± SEM from 4 wild-type male mouse hypothalami, with 12 to 15 sections per animal. Scale bars: panoramic views, 200 µm; high-magnification images, 20 µm.

### 3.2. 26RFa and orexin exert convergent actions on glucose homeostasis

Given their extensive co-expression within the same lateral hypothalamic neurons, we next asked whether 26RFa and orexin exert comparable actions on glucose homeostasis. To address this question, we combined complementary genetic and pharmacological approaches.

We first examined whether orexin deficiency phenocopies the metabolic alterations previously reported in 26RFa-deficient mice and whether combined deficiency of both neuropeptides further aggravates glucose dysregulation. Body weight measurement from postnatal week 5 to week 13 indicated that the orexin-deficient mice and the double 26RFa/orexin deficient mice were significantly heavier than wild-type controls (suppl Fig. 2A; Fig. 2A). By contrast, basal plasma glucose levels measured after a 16-h fasting did not differ significantly between the two groups (suppl Fig. 2B; Fig. 2B). An intraperitoneal glucose tolerance test revealed a marked increase in the hyperglycemic peak induced by glucose loading, in orexin-deficient and double 26RFa/orexin deficient mice (suppl Fig. 2C; Fig. 2C). An insulin tolerance test indicated that orexin deficiency alone or associated with 26RFa deficiency is accompanied by a slight decrease in insulin sensitivity throughout the test (suppl Fig. 2D; Fig. 2D). Acute glucose-stimulated insulin secretion was decreased in both orexin-and 26RFa/orexin-deficient mice with a more pronounced effect observed in the double KO mice (suppl Fig. 2E; Fig. 2E). Finally, a pyruvate tolerance test showed a sustained increase of the hepatic glucose production over the course of the test in orexin-deficient mice and in mice deficient in both 26RFa and orexin (suppl Fig. 2F; Fig. 2F). Together, these results indicate that orexin deficiency reproduces key metabolic defects previously described in 26RFa-deficient mice and that combined deficiency further aggravates glucose homeostasis, supporting complementary contributions of the two neuropeptidergic systems.

**Figure 2.**
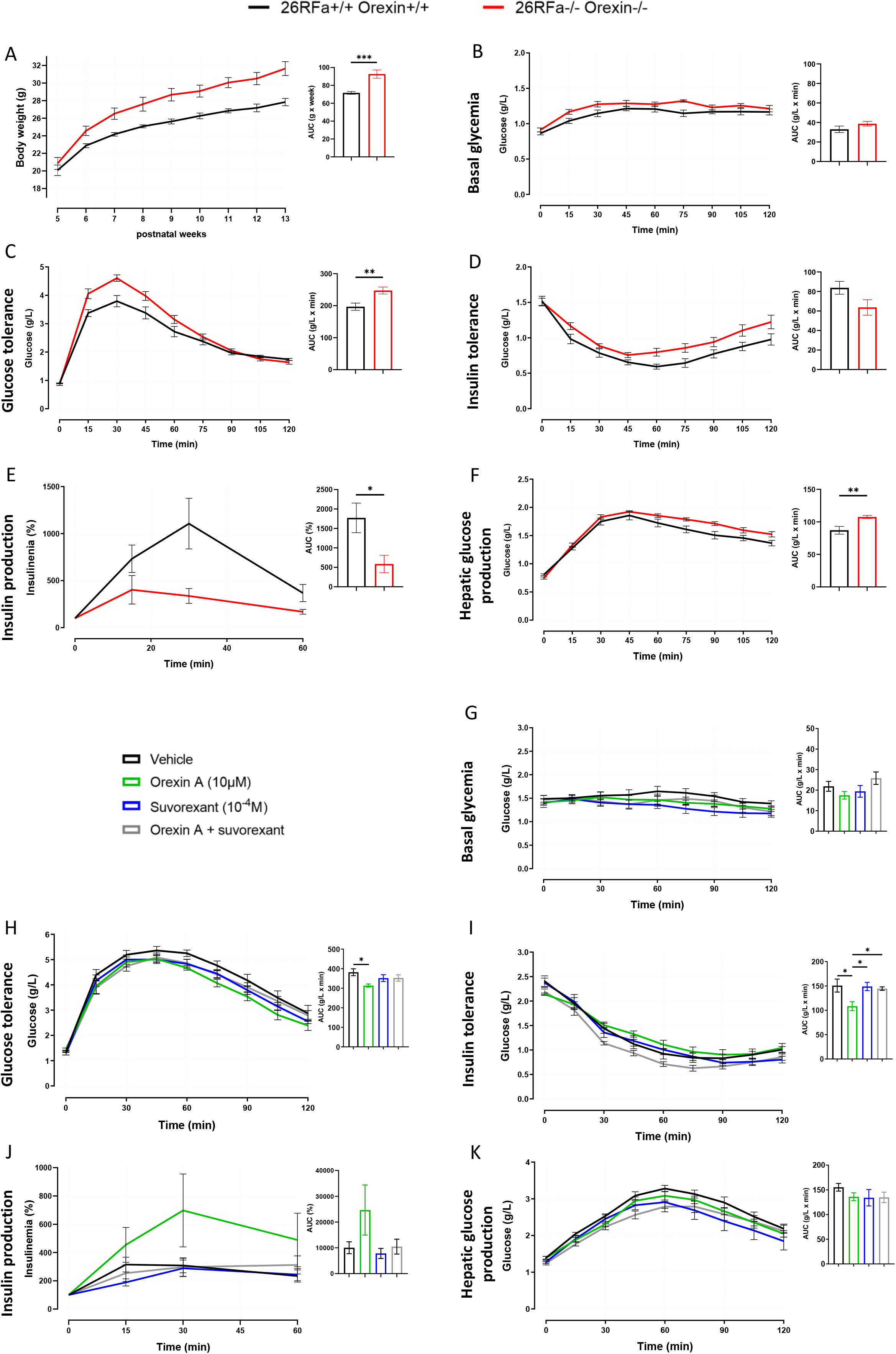
26RFa and orexin exert convergent actions on glucose homeostasis. **A)** Body weight evolution from postnatal week 5 to 13 in 26RFa^-/-^;Orexin^-/-^mice (red curve) compared to wild-type animals (black curve). **B, C, D, F)** Impact of double 26RFa/orexin deficiency (red curves) on plasma glucose levels under basal conditions (**B**), during an intraperitoneal (i.p.) glucose tolerance test (**C**), an insulin tolerance test (**D**), and an i.p. pyruvate tolerance test (**F**). **E)** Effect of double 26RFa/orexin deficiency (red curve) on plasma insulin levels during the i.p. glucose tolerance test. **G, H, I, K)** Effect of intracerebroventricular (i.c.v.) administration of orexin A (red curves), or the orexin receptor antagonist suvorexant (blue curves) or their combination (grey curves), or vehicle (black curves) on plasma glucose levels under basal conditions (**G**), during an i.p. glucose tolerance test (**H**), an insulin tolerance test (**I**), and an i.p. pyruvate tolerance test (**K**). **J)** Effect of i.c.v. administration of orexin A (red curve), suvorexant (blue curve), or orexin A + suvorexant (grey curve) or vehicle (black curve) on plasma insulin levels during the i.p. glucose tolerance test. Graphs and AUC values are represented as mean ± SEM of 10 mice per group. *p < 0.05 ; \*\**p*<0.01; \*\*\**p*<0.001.

To determine whether orexin signaling itself exerts central metabolic actions comparable to those previously described for 26RFa, we next assessed the effects of central orexin administration. Neither central administration of orexin A, nor the orexin receptor antagonist suvorexant affect basal plasma glucose levels (Fig. 2G). By contrast, central orexin A significantly attenuates glucose-induced hyperglycemia during a glucose tolerance test (Fig. 2H), and this effect is blocked by co-administration of suvorexant, whereas suvorexant alone has no effect on plasma glucose levels (Fig. 2H). An insulin tolerance test showed that orexin A significantly attenuates the hypoglycemic effect of insulin, an effect reversed by the receptor antagonist (Fig. 2I). An acute glucose-stimulated insulin secretion (AGSIS) test revealed that i.c.v. injection of orexin A potentiates glucose-induced insulin production, an effect abolished by the co-administration of the orexin receptor antagonist (Fig. 2J). Finally, pyruvate tolerance testing indicated that central orexin A does not significantly affect the hepatic glucose production (Fig. 2K). Together, these complementary genetic and pharmacological approaches demonstrate that orexin, like 26RFa, exerts predominantly anti-hyperglycemic actions at the central level, supporting convergent functions of the two peptidergic systems within the newly identified 26RFa/orexin neuronal population.

### 3.3. Molecular relationship between the 26RFa and orexin peptidergic systems

Because 26RFa and orexin exert convergent metabolic actions within the same neuronal population, we next asked whether these two peptidergic systems are functionally linked at the molecular level. In 26RFa-deficient mice, RNAscope analyses revealed a marked decrease in both the number and labelling intensity of orexin-expressing neurons, whereas the number of GPR103-positive neurons remained unchanged (Fig. 3A). These findings indicate that loss of 26RFa selectively remodels the orexin component of this neuronal population while preserving GPR103 expression. To further examine this relationship, we assessed the impact of central pharmacological modulation of 26RFa signaling on orexin expression. i.c.v. administration of 26RFa under hyperglycemic conditions tends to increase orexin mRNA expression in the LHA, whereas co-administration with the GPR103 antagonist, 25e, significantly reduces orexin expression (Fig. 3B), supporting a positive regulatory influence of 26RFa signaling on orexin transcriptional control.

**Figure 3.**
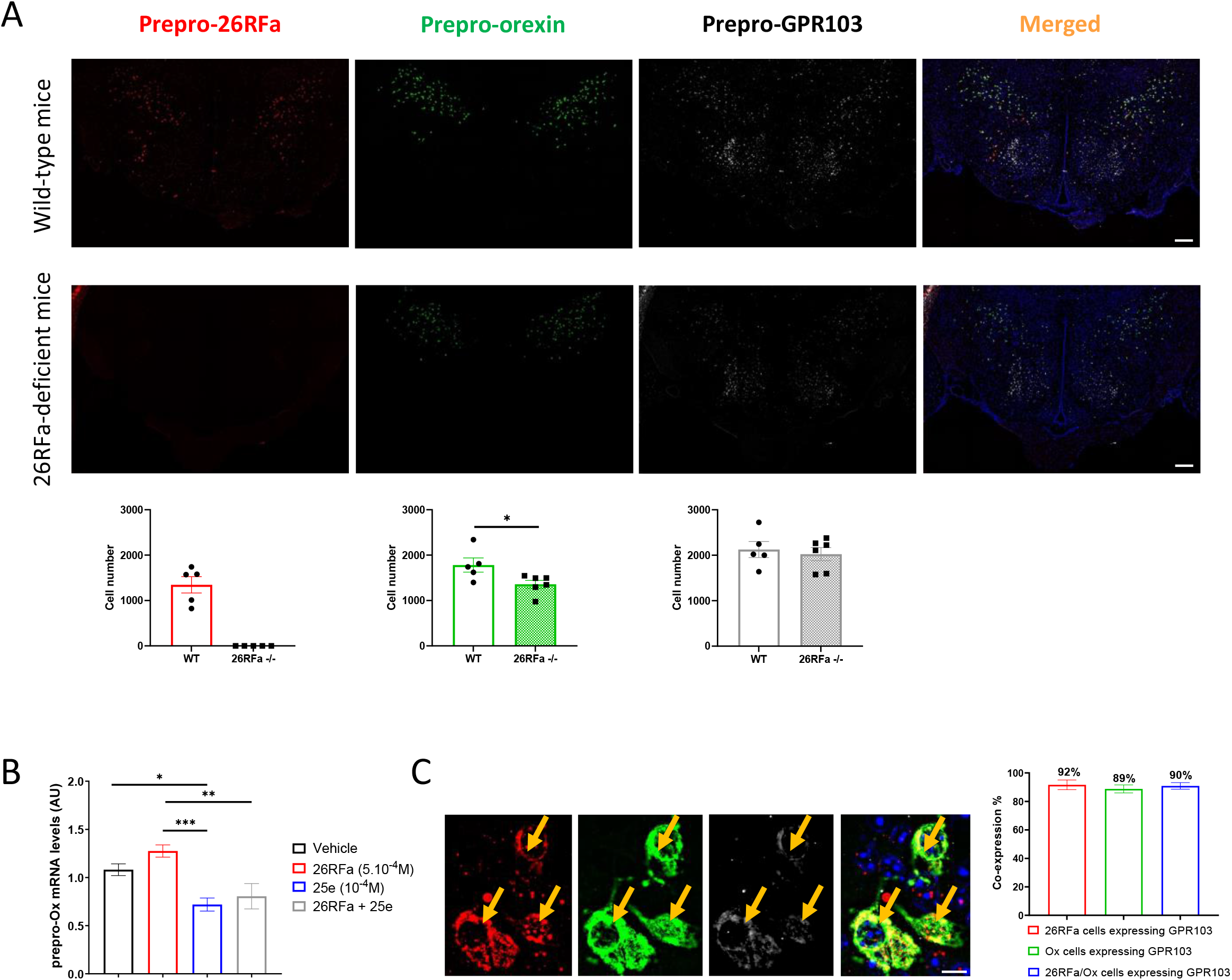
26RFa signaling positively regulates orexin expression within the LHA. **A)** Panoramic views of hypothalamic sections showing the distribution of *prepro-26RFa* (red), *prepro-orexin* (green) and *prepro-GPR103* (grey) mRNAs, as revealed by RNAscope *in situ* hybridization in wild-type and 26RFa-deficient mice. Graphs represent the impact of 26RFa deficiency on the number of 26RFa-expressing cells (red), orexin-expressing cells (green) and GPR103-expressing cells (grey). **B)** Effect of intracerebroventricular administration of vehicle (black), 26RFa (red), the GPR103 antagonist 25e (blue) and 26RFa + 25e (grey) on the hypothalamic *prepro-orexin* expression. **C)** Representative photomicrographs of cells in the lateral hypothalamic area co-expressing *prepro-26RFa*, *prepro-orexin* and *prepro-GPR103* (yellow arrow), along with the quantitative analysis of co-expressing cells. Data are presented as mean ± SEM of hypothalami from 6 26RFa^+/+^ male mice, with 10–12 sections analyzed per animal. Scale bars: panoramic views, 200 µm; high-magnification images, 20 µm. \**p*<0.05; \*\**p*<0.01; \*\*\**p*<0.001.

Finally, RNAscope® *in situ* hybridization revealed that the vast majority (∼90%) of 26RFa/orexin-expressing neurons co-express the GPR103 in the LHA (Fig. 3C), indicating that most neurons within this population are equipped to directly respond to 26RFa signaling. Together, these findings indicate that the 26RFa and orexin peptidergic systems are molecularly coupled within the same neuronal population of the LHA, with 26RFa positively regulating orexin expression and GPR103 representing a potential mediator of this interaction.

### 3.4. Neuronal activity coordinates glycemic and transcriptional responses within LHA 26RFa/orexin neurons

Having established that 26RFa and orexin form a molecularly coupled peptidergic system within the same neuronal population, we next investigated how changes in neuronal activity influence both systemic glucose homeostasis and transcriptional regulation of these two neuropeptidergic systems. To this end, LHA 26RFa/orexin neurons were selectively activated or inhibited using a chemogenetic approach in 26RFa/orexin-expressing mice.

We first chemogenetically activated this neuronal population by expressing excitatory DREADDs specifically under the control of the 26RFa promoter. Control experiments using empty AAV vectors injected demonstrated that CNO administration does not affect glycemic parameters, except for a modest pro-hyperglycemic effect during glucose tolerance testing (Suppl. Fig. 3). Chemogenetic activation of the LHA 26RFa/orexin neurons does not affect basal glycemia (Fig. 4A) and insulin sensitivity (Fig. 4E). By contrast, activation of this neuronal population significantly potentiates glucose-induced hyperglycemia throughout a glucose tolerance test (Fig. 4B) accompanied by a marked increase in glucose-stimulated insulin secretion (Fig. 4C) and a modest increase in hepatic glucose production detected by pyruvate tolerance testing (Fig. 4D.

**Figure 4.**
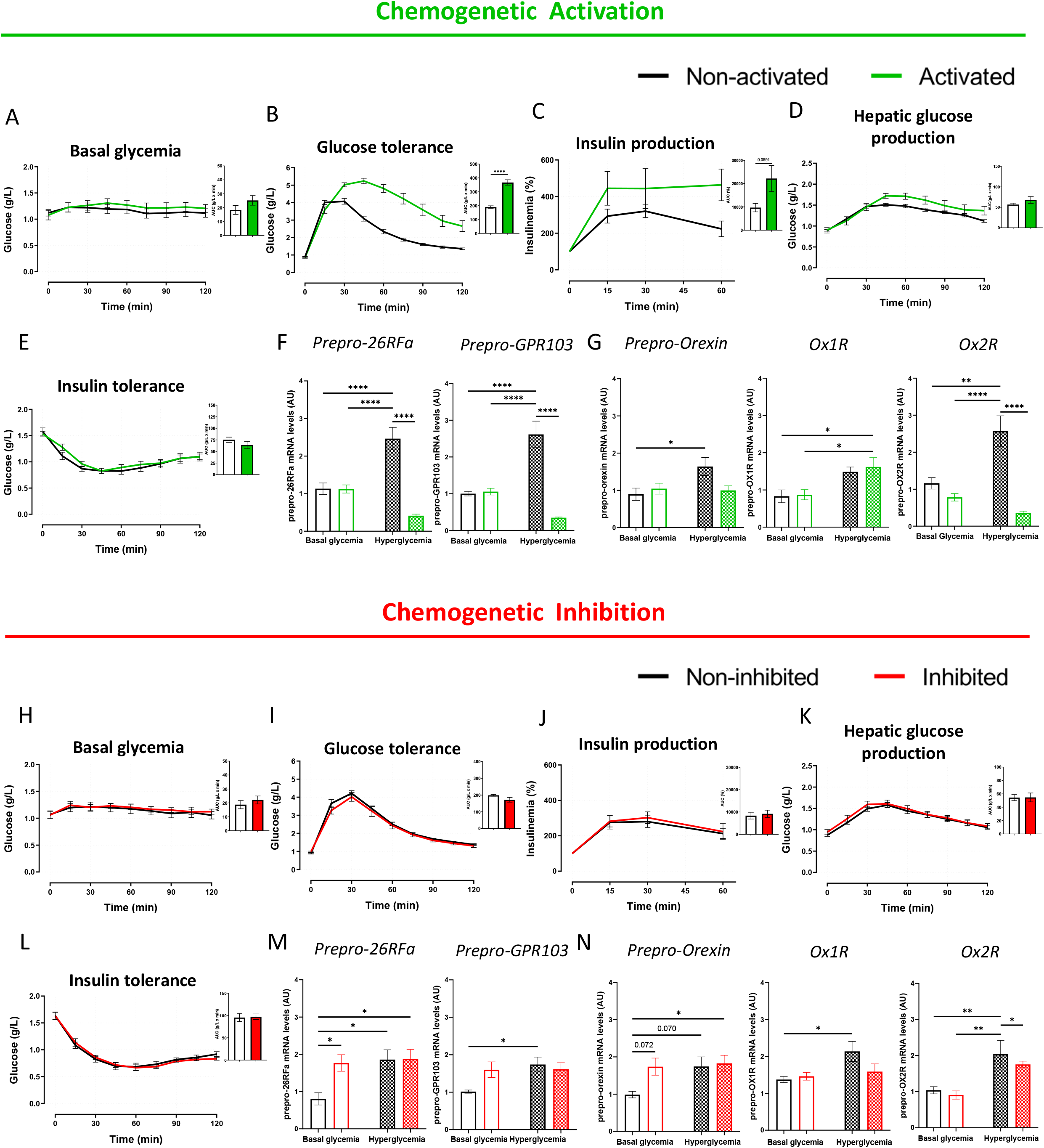
Activity-dependent regulation of the 26RFa and orexin peptidergic systems in wild-type mice. **A-E)** Impact of chemogenetic activation of LHA 26RFa neurons (green curves), compared to control condition (black curves), on basal glycemia **(A)**, glucose tolerance **(B)**, insulin production **(C)**, hepatic glucose production **(D)** and insulin tolerance **(E)** in wild-type mice. **F-G)** Effect of chemogenetic activation of 26RFa/orexins neurons on the hypothalamic expression of *prepro-26RFa*/*prepro-GPR103* **(F)** and of *prepro*-*orexin*/*Ox1R*/*Ox2R* **(G)** under basal glycemic conditions (16 h fasting mice) or under hyperglycemic conditions (45min after an i.p. glucose challenge in 16 h fasting mice) in wild-type mice. **H-L)** Impact of chemogenetic inhibition of LHA 26RFa neurons (red curves), compared to control condition (black curves), on basal glycemia **(H)**, glucose tolerance **(I)**, insulin production **(J)**, hepatic glucose production **(K)** and insulin tolerance **(L)** in wild-type mice. **M-N)** Effect of chemogenetic inhibition of 26RFa/orexins neurons on the hypothalamic expression of *prepro-26RFa/prepro-GPR103* **(M)** and of *prepro*-*orexin*/*Ox1R*/*Ox2R* **(N)** in basal glycemic condition (16 h fasting mice) or in hyperglycemic condition (45 min after an i.p. glucose challenge in 16 h fasting mice) in wild-type mice. All graphs and corresponding AUC are presented as mean ± SEM of 8-10 mice (non-activated vs activated) with each mouse serving as its own control. \**p*<0.05; \*\**p*<0.01; \*\*\**p*<0.001; \*\*\*\**p*<0.0001.

This paradoxical pro-hyperglycemic response, elicited by activation of neurons co-expressing two anti-hyperglycemic neuropeptides, prompted us to investigate whether neuronal activation was associated with transcriptional adaptations of the two peptidergic systems, under different glycemic conditions. Under physiological conditions, systemic hyperglycemia induces a coordinated upregulation of both 26RFa and orexin peptidergic systems, including increased expression of 26RFa, GPR103, orexin and the two orexin receptors (Fig. 4F, G). We therefore examined whether chemogenetic activation modified this physiological transcriptional response. Under basal glycemic conditions, corresponding to the mild hypoglycemia induced by a 16-h fasting period, activation of LHA 26RFa neurons does not alter the expression of either peptidergic system (Fig. 4F, G). In contrast, under hyperglycemic conditions, neuronal activation reverses the transcriptional response, resulting in a coordinated transcriptional downregulation of 26RFa, GPR103, orexin and Ox2R expression (Fig. 4F, G). Together, these findings indicate that chemogenetic activation of LHA 26RFa/orexin neurons is associated with both potentiation of glucose-induced hyperglycemia and coordinated transcriptional repression of the 26RFa and orexin peptidergic systems.

We next examined the consequences of chemogenetic inhibition of LHA 26RFa/orexin neurons. In contrast to neuronal activation, inhibition did not affect basal glycemia, glucose tolerance, glucose-stimulated insulin secretion, hepatic glucose production or insulin sensitivity (Fig. 4H–L), indicating that reduced neuronal activity has only limited consequences on systemic glucose homeostasis under these conditions. At the transcriptional level, chemogenetic inhibition produced a distinct pattern depending on glycemic status. Under basal glycemic conditions, inhibition significantly increases hypothalamic 26RFa expression (Fig. 4M), with concomitant trends toward increased GPR103 and orexin expression, whereas orexin receptor levels remain unchanged (Fig. 4M, N). Under hyperglycemic conditions, inhibition of these neurons has no impact on the expression of the 26RFa and orexin peptidergic systems of the hypothalamus except for a slight decrease in Ox2R expression (Fig. 4M, N). These findings indicate that, unlike neuronal activation, chemogenetic inhibition exerts only limited effects on glucose homeostasis and does not substantially modify the physiological transcriptional response to hyperglycemia, while selectively increasing 26RFa expression under basal glycemic conditions.

Together, these findings reveal that the molecularly coupled 26RFa and orexin peptidergic systems are dynamically regulated by neuronal activity in a glycemic state-dependent manner. While chemogenetic activation overrides the physiological transcriptional response to hyperglycemia and is associated with paradoxical potentiation of glucose-induced hyperglycemia, neuronal inhibition primarily influences transcription under basal glycemic conditions with minimal glycemic consequences.

### 3.5. Role of 26RFa signaling in the activity-dependent organization of LHA 26RFa/orexin neurons

We then examined whether the activity-dependent organization identified in wild-type mice depended on intact 26RFa signaling. To address this question, we performed chemogenetic activation and inhibition of LHA 26RFa/orexin neurons in 26RFa-deficient mice.

Overall, chemogenetic activation produced glycemic effects largely comparable to those observed in wild-type mice. Basal glycemia remains unchanged, whereas glucose-induced hyperglycemia is still potentiated during a glucose tolerance test (Fig. 5A, B). However, two notable differences emerge. First, the increase in glucose-stimulated insulin secretion observed in wild-type mice is completely lost in 26RFa-deficient animals (Fig. 5C). Second, hepatic glucose production is further enhanced following neuronal activation (Fig. 5D), and insulin sensitivity is reduced (Fig. 5E). Unlike wild-type mice, chemogenetic inhibition of LHA 26RFa/orexin neurons now induces a modest but significant alteration of glucose tolerance (Fig. 5I), whereas basal glycemia, insulin secretion, hepatic glucose production and insulin sensitivity remain unaffected (Fig. 5H, J-L).

**Figure 5.**
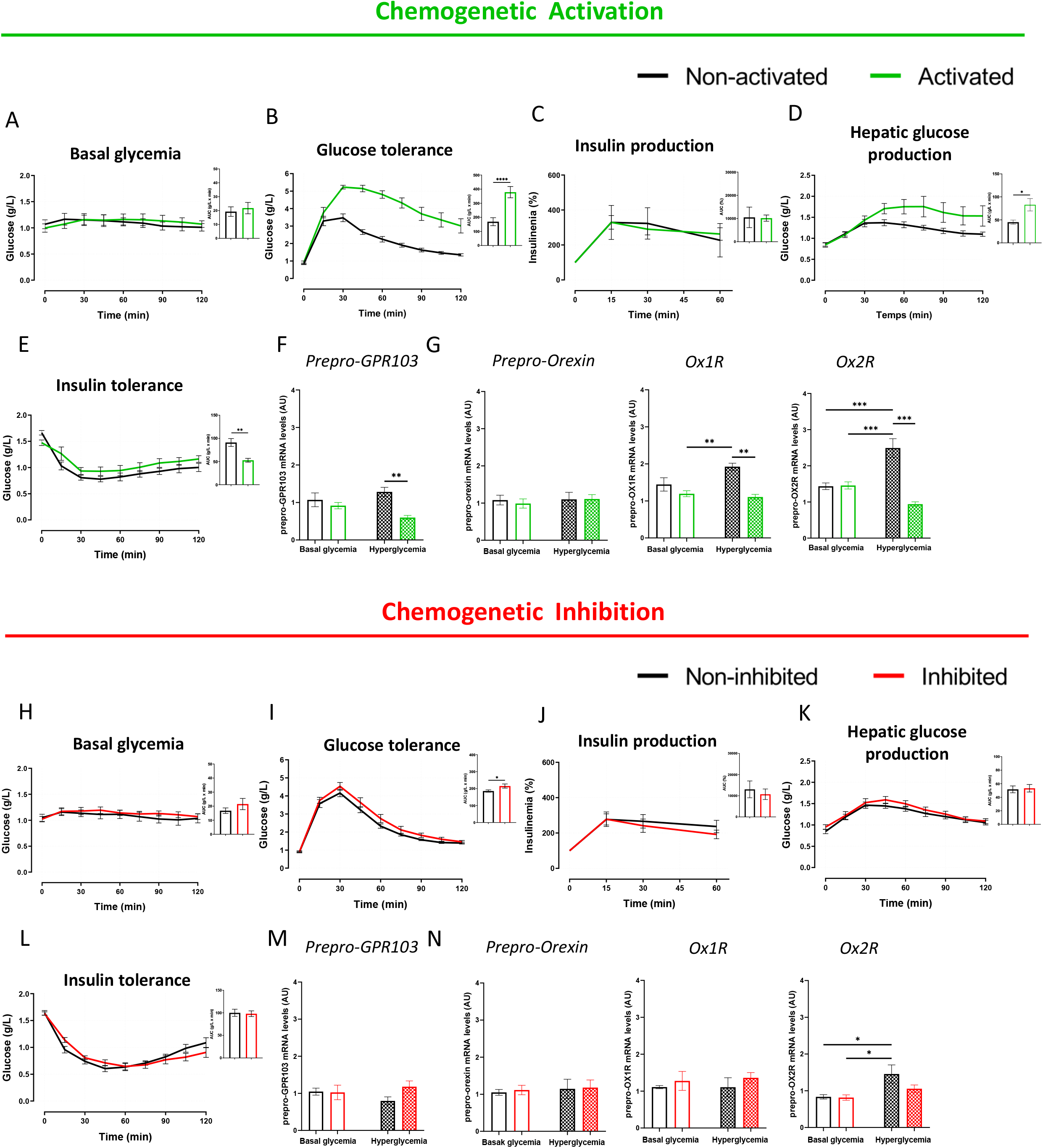
Loss of 26RFa signaling disrupts the activity-and glycemic state-dependent regulation of the orexin peptidergic system. **A-E)** Impact of chemogenetic activation of LHA 26RFa neurons (green curves), compared to control condition (black curves), on basal glycemia **(A)**, glucose tolerance **(B)**, insulin production **(C)**, hepatic glucose production **(D)** and insulin tolerance **(E)** in 26RFa-deficient mice. **F-G)** Effect of chemogenetic activation of 26RFa/orexins neurons on the hypothalamic expression of *prepro-GPR103* **(F)** and of *prepro*-*orexin*/*Ox1R*/*Ox2R* **(G)** in basal glycemic condition (16 h fasting mice) or in hyperglycemic condition (45min after an i.p. glucose challenge in 16 h fasting mice) in 26RFa-deficient mice. **H-L)** Impact of chemogenetic inhibition of LHA 26RFa neurons (red curves), compared to control condition (black curves), on basal glycemia **(H)**, glucose tolerance **(I)**, insulin production **(J)**, hepatic glucose production **(K)** and insulin tolerance **(L)** in 26RFa-deficient mice. **M-N)** Effect of chemogenetic inhibition of 26RFa/orexins neurons on the hypothalamic expression of *prepro-GPR103* **(M)** and of *prepro*-*orexin*/*Ox1R*/*Ox2R* **(N)** under basal glycemic conditions (16 h fasting mice) or under hyperglycemic conditions (45 min after an i.p. glucose challenge in 16 h fasting mice) in 26RFa-deficient mice. All graphs and corresponding AUC are mean ± SEM of 8-10 mice (non-activated vs activated) with each mouse serving as its own control. \**p*<0.05; \*\**p*<0.01; \*\*\**p*<0.001; \*\*\*\**p*<0.0001.

In wild-type mice, systemic hyperglycemia induced coordinated upregulation of both the 26RFa and orexin peptidergic systems (Fig. 4F, G). We therefore asked whether this physiological transcriptional response is preserved in the absence of 26RFa signaling. Unlike wild-type mice, 26RFa-deficient animals fail to exhibit this glycemic state-dependent transcriptional response. Indeed, systemic hyperglycemia no longer increases GPR103, orexin or Ox1R expression, whereas Ox2R expression is selectively increased (Fig. 5F-G, M-N). Chemogenetic activation produces a transcriptional profile that is also altered by the loss of 26RFa signaling. As observed in wild-type mice, activation under basal glycemic conditions does not modify the expression of GPR103, orexin or orexin receptors (Fig. 5F, G). Under hyperglycemic conditions, activation still reduces GPR103 and orexin receptor expression, whereas orexin remains unchanged, consistent with the absence of glucose-induced orexin upregulation in 26RFa-deficient mice (Fig. 5F, G). The transcriptional response to chemogenetic inhibition is likewise altered. Whereas inhibition increases 26RFa expression under basal glycemic conditions in wild-type mice, it no longer modifies GPR103, orexin or orexin receptor expression in 26RFa-deficient animals (Fig. 5M, N). Similarly, inhibition fails to alter transcription under hyperglycemic conditions, except for the modest reduction in Ox2R expression already observed in wild-type mice (Fig. 5M, N). These findings reveal that loss of 26RFa signaling profoundly alters the activity- and glycemic state-dependent transcriptional regulation of the 26RFa and orexin peptidergic systems.

### 3.6. Role of orexin signaling in the activity-dependent organization of LHA 26RFa/orexin neurons

We next assessed the contribution of orexin signaling to the glycemic responses elicited by chemogenetic modulation of LHA 26RFa/orexin neurons. Basal glycemia remained unaffected by either activation or inhibition in orexin-deficient and double-deficient mice (Fig. 6A–F). In orexin-deficient mice, chemogenetic activation still potentiated glucose-induced hyperglycemia, although this response was markedly attenuated compared with wild-type and 26RFa-deficient mice (Fig. 6G, I). This effect was completely abolished in double-deficient mice (Fig. 6H, I). Likewise, the activation-induced increase in insulin secretion observed in wild-type mice was absent in both orexin-deficient and double-deficient animals (Fig. 6M–O). Similarly, activation no longer increased hepatic glucose production in either genotype (Fig. 6S–U), while insulin sensitivity remained unchanged (Fig. 6Y–AA).

**Figure 6.**
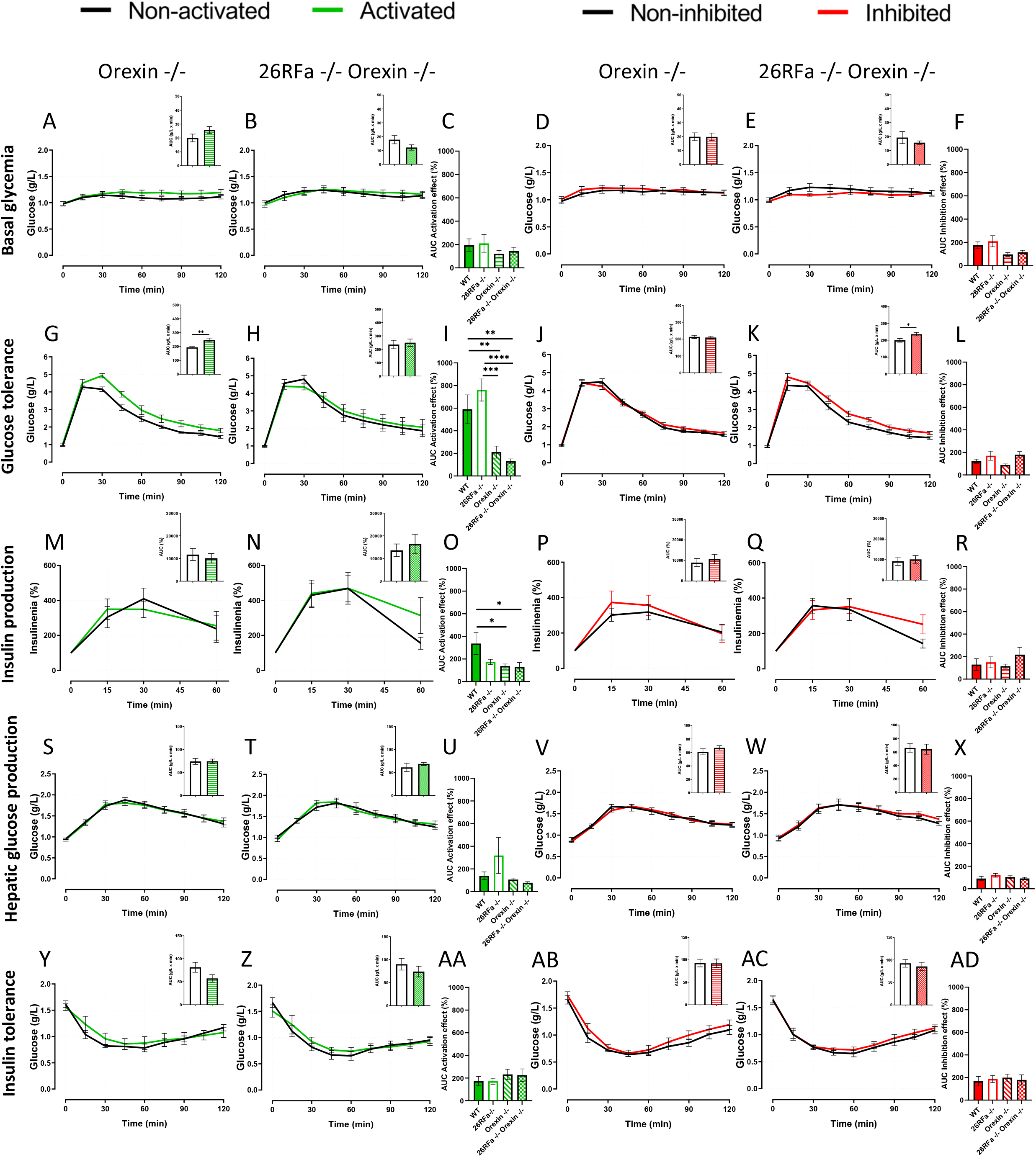
Orexin signaling is required for the full glycemic response induced by neuronal activation. **A–B, G–H, M–N, S–T, Y–Z)** Impact of chemogenetic activation of LHA 26RFa neurons (green curves) compared to control condition (black curves), on basal glycemia **(A, B,)**, glucose tolerance **(G, H)**, insulin production **(M, N)**, hepatic glucose production **(S, T)** and insulin tolerance **(Y, Z)** in orexin-deficient mice **(A, G, M, S, Y)** and in double 26RFa/orexin deficient **mice (B, H, N, T, Z)**. **D–E, J–K, P–Q, V–W, AB–AC)**: Impact of chemogenetic inhibition of LHA 26RFa neurons (red curves) compared to control condition (black curves), on basal glycemia **(D, E)**, glucose tolerance **(J, K)**, insulin production **(P, Q)**, hepatic glucose production **(V, W)** and insulin tolerance **(AB, AC)** in orexin-deficient mice **(D, J, P, V, AB)** and in double 26RFa/orexin deficient **mice (E, K, Q, W, AC)**. **C, I, O, U, AA)** Area under the curve (AUC) values representing the effect of chemogenetic activation (%). **F, L, R, X, AD):** AUC values representing the effect of chemogenetic inhibition (%). To isolate the specific effect of chemogentic modulation, glycemia values measured after CNO administration at each time point were normalized to the corresponding values obtained under the control condition. This normalization yielded a relative chemogenetic modulation curve (% of control), from which the AUC was calculated to represent the overall chemogenetic modulation effect. Data are presented as mean ± SEM of 8-10 mice, with each mouse serving as its own control. For normalized curves, values obtained following chemogenetic modulation are expressed as a percentage of the corresponding control values.. \**p*<0.05; \*\**p*<0.01; \*\*\**p*<0.001; \*\*\*\**p*<0.0001.

Chemogenetic inhibition produced only limited glycemic effects in orexin-deficient mice and failed to modify insulin secretion, hepatic glucose production or insulin sensitivity (Fig. 6P–R, V–X, AB–AD). Similarly, inhibition remained without effect in double-deficient mice, except for the modest impairment in glucose tolerance previously associated with the loss of 26RFa signaling (Fig. 6J–L).

Collectively, these findings indicate that orexin signaling is required for the full expression of the metabolic responses elicited by activation of LHA 26RFa/orexin neurons, particularly the pro-hyperglycemic response and its associated insulin and hepatic components.

## 4. Discussion

Hypothalamic circuits that sense metabolic signals play a central role in the regulation of systemic glucose homeostasis. While much attention has focused on glucose-responsive networks in the Arc, the contribution of the lateral hypothalamus to central glucose regulation remains less clearly defined despite its established role in metabolic control. Here, we investigate the functional organization of a population of lateral hypothalamic neurons co-expressing the neuropeptides 26RFa and orexin and assess their contribution to the central regulation of glucose homeostasis. Surprisingly, chemogenetic activation of these neurons produces a pro-hyperglycemic response despite the anti-hyperglycemic effects of both peptides when administered centrally. This paradoxical response is associated with coordinated, activity- and glycemic state-dependent transcriptional regulation of both the 26RFa and orexin peptidergic systems. Together, these findings identify a close relationship between neuronal activity, transcriptional regulation and systemic glucose homeostasis within LHA 26RFa/orexin neurons.

Neuroanatomical studies previously identified a population of 26RFa-expressing cells within the LHA (El Mehdi *et al*, 2022; Seifinejad *et al*, 2023; Picot *et al*, 2025), a hypothalamic region classically recognized for its role in the control of energy balance and metabolic homeostasis. The LHA also contains two major peptidergic neuronal populations, the orexin- and melanin-concentrating hormone (MCH)-producing neurons (Bonnavion *et al*, 2016), both of which have been suspected to contribute to the regulation of glucose homeostasis (Tsuneki *et al*, 2012; Cook *et al*, 2022). Despite this metabolic relevance, the cellular identity and organization of LHA 26RFa-expressing cells have remained incompletely defined. Here, using complementary neuroanatomical and molecular approaches, we provide a detailed characterization of this population and show that 26RFa-expressing cells in the LHA display extensive co-expression with orexin (∼90%), while approximately two-thirds of these cells exhibit a neuronal identity (∼66%). These findings support very recent observations of 26RFa-orexin co-expression (Seifinejad *et al*, 2023) and identify a discrete lateral hypothalamic peptidergic population within the LHA, referred to here as 26RFa/orexin neurons. The presence of a fraction of non-neuronal 26RFa-expressing cells suggests additional cellular complexity within this hypothalamic population that deserves further investigation.

The functional significance of this dual peptidergic identity is notable in light of the metabolic actions of the two peptides. Previous work demonstrated that central administration of 26RFa reduces glucose-induced hyperglycemia by stimulating pancreatic insulin secretion and that the hypothalamic 26RFa/GPR103 system acts as a relay of central insulin signaling (El Mehdi *et al*, 2022). In the present study, we show that central administration of orexin A produces closely related metabolic effects, including an anti-hyperglycemic response associated with increased insulin production and decreased insulin sensitivity. Thus, 26RFa/orexin neurons contain two neuropeptides that exert convergent central actions on glucose regulation. The coexistence of these functionally similar peptidergic signals within the same neuronal population raises the question of how these two peptidergic systems are coordinated within the lateral hypothalamic circuitry controlling glucose homeostasis.

The co-expression of 26RFa and orexin, together with the convergence of their metabolic actions, prompted us to investigate how modulation of neuronal activity influences systemic glucose homeostasis. Strikingly, chemogenetic activation of LHA 26RFa/orexin neurons enhances the hyperglycemic response to a peripheral glucose load while simultaneously stimulating pancreatic insulin secretion. Notably, activation has no detectable effect on basal glycemia, insulin sensitivity, or hepatic glucose production. This paradoxical pro-hyperglycemic effect was unexpected, as 26RFa has been previously shown to exert robust anti-hyperglycemic effects upon central administration (El Mehdi *et al*, 2022), and we show here that orexin produces a similar effect. Interestingly, a recent study shows that chemogenetic activation of 26RFa-expressing neurons in the preoptic area similarly induces a pro-hyperglycemic response during a glucose tolerance test, also associated with increased insulin production (Kong *et al*, 2010). In line with these findings, LHA orexin neurons are known to be activated under hypoglycemic conditions (Paranjape *et al*, 2006) and inhibited in hyperglycemic environments (Burdakov & González, 2009; Sheng *et al*, 2014), supporting the idea that artificial activation of 26RFa/orexin neurons may mimic physiological responses normally engaged under hypoglycemic conditions, thereby contributing to the paradoxical increase in glycemia.

To understand the basis of this apparent paradox, we assessed transcriptional changes in 26RFa/orexin neurons. Hypothalamic RT-qPCR analyses revealed that hyperglycemia upregulates both 26RFa and orexin mRNA expression, whereas chemogenetic activation of these neurons abolishes this transcriptional response. Conversely, their chemogenetic inhibition under basal glycemic conditions (mildly hypoglycemic after a 16 h fast) results in an upregulation of 26RFa and orexin transcripts to levels comparable to those observed in hyperglycemia. Taken together with previous studies, our findings support the idea that LHA 26RFa/orexin neurons are preferentially active under basal and mild hypoglycemic conditions and become progressively inhibited as glycemia rises. We therefore propose that coordinated downregulation of the 26RFa and orexin peptidergic systems represents a candidate counter-regulatory mechanism that may contribute to the paradoxical pro-hyperglycemic response elicited by sustained neuronal activation.

To dissect the respective contributions of 26RFa and orexin to glucose homeostasis and to the paradoxical response induced by neuronal activation, we examined mice deficient for each peptide individually as well as double mutants lacking both neuropeptides. Double knockout mice develop severe obesity and exhibit a markedly impaired glucose tolerance, associated with impaired insulin secretion, mildly reduced insulin sensitivity, and increased hepatic glucose production. This phenotype is more pronounced than that observed in either 26RFa-or orexin-deficient mice (El Mehdi *et al*, 2022; Devère *et al*, 2025), suggesting additive or synergistic contributions of the two peptidergic systems to glucose homeostasis.

We next examined how chemogenetic activation of 26RFa/orexin neurons impacts glucose tolerance among these genetic backgrounds. The pro-hyperglycemic response to neuronal activation is preserved in 26RFa-deficient mice, markedly attenuated in orexin-deficient mice, and completely abolished in double knockout animals. The attenuation observed in orexin-deficient mice likely reflects an elevated basal glycemic state, which limits the dynamic range of further increases during glucose challenge. Consistent with this interpretation, the absence of any glycemic response in double knockout mice can be explained by their severely impaired glucose tolerance at baseline. These observations indicate that both neuropeptides contribute to shaping the glycemic response.

Despite these differences in glycemic outcomes, the activation-induced increase in insulin secretion observed in wild-type mice is lost in all mutant strains, including 26RFa-, orexin-, and double-deficient mice, indicating that the insulinotropic component strictly depends on the integrity of both peptidergic systems, consistent with previous reports showing impaired glucose-stimulated insulin secretion in both 26RFa- and orexin-deficient mice (Devère *et al*, 2025; El Mehdi *et al*, 2020) and in double-deficient mice, whereas the overall glycemic outcome reflects additional regulatory mechanisms. Consistent with this dissociation, the increase in hepatic glucose production is modest and does not reach statistical significance in wild-type mice, whereas it becomes significant in 26RFa-deficient mice following neuronal activation. By contrast, no effect is observed in orexin- or double-deficient mice. This pattern indicates that the activation-dependent modulation of hepatic glucose production requires orexin signaling. Notably, although both 26RFa- and orexin-deficient mice display elevated basal hepatic glucose production, only 26RFa-deficient mice retain the capacity to further increase this parameter upon neuronal activation, whereas this dynamic response is lost in orexin- and double-deficient animals. Together, these findings indicate that LHA 26RFa/orexin neurons contribute to the regulation of systemic glucose homeostasis in an activity-dependent manner, with neuronal activation paradoxically promoting hyperglycemia while being associated with coordinated downregulation of both peptidergic systems. In this context, both peptidergic systems are required for the insulinotropic component of the response, whereas orexin signaling preferentially contributes to the regulation of hepatic glucose production, consistent with a modulatory role of 26RFa on this pathway under physiological conditions.

To further assess the functional relevance of this activity-dependent regulation, we examined the effects of chemogenetic inhibition of LHA 26RFa/orexin neurons on glucose homeostasis. In contrast to neuronal activation, inhibition of this population produces only limited effects on glycemic control and does not alter basal glycemia, insulin secretion, insulin sensitivity, or hepatic glucose production among the different genotypes. Notably, inhibition induces a modest but significant pro-hyperglycemic effect selectively in mice lacking 26RFa signaling, including both 26RFa-deficient and double-deficient animals, whereas minimal effects are observed in wild-type and orexin-deficient mice. Although no significant effect is detected in wild-type mice, control experiments indicating a mild pro-hyperglycemic effect of CNO suggest that a potential inhibitory component may be partially masked under these conditions. Interestingly, this intrinsic pro-hyperglycemic effect of CNO is no longer observed when 26RFa/orexin neurons are inhibited in mice expressing both neuropeptides, suggesting that neuronal inhibition counteracts the effect of CNO under these conditions.

Chemogenetic inhibition in wild-type mice increases the expression of both 26RFa and orexin, yet this transcriptional upregulation does not translate into an anti-hyperglycemic effect. We propose that this apparent dissociation reflects the fact that expression of both neuropeptides is already elevated under basal or mild hypoglycemic conditions, thereby limiting the impact of further transcriptional upregulation during glucose challenge. Together, these observations highlight a functional asymmetry within the 26RFa/orexin system, in which neuronal activation and inhibition do not produce mirror effects but instead reveal distinct functional contributions of the two peptidergic systems. We propose that orexin signaling primarily mediates the metabolic response to neuronal activation, whereas 26RFa acts as a modulatory component that shapes this response. Within this framework, the two neuropeptidergic systems appear functionally complementary rather than redundant.

The distinct functional contributions of 26RFa and orexin signaling suggest that the two peptidergic systems are not simply co-expressed but are functionally coordinated within LHA neurons. Our molecular analyses support this view as, in intact circuits, both neuropeptidergic systems display coordinated glycemic state-dependent transcriptional regulation, with hyperglycemia inducing their upregulation and neuronal activation under hyperglycemic conditions leading to their coordinated repression. This coupling is disrupted in 26RFa- deficient mice, in which glycemic state-dependent transcriptional modulation of the orexin system is lost, suggesting that 26RFa signaling contributes to the dynamic regulation of orexin expression. Consistent with this view, chemogenetic inhibition or hyperglycemia fail to induce orexin upregulation in the absence of 26RFa, and RNAscope analyses reveal a reduction in both the number and labeling intensity of orexin-expressing neurons in 26RFa-deficient mice. Pharmacological manipulations further support this relationship, as blockade of the 26RFa receptor reduces orexin expression under hyperglycemic conditions, whereas central administration of 26RFa has been reported to increase it (Lee *et al*, 2025). Together with previous reports showing that orexin deficiency does not affect 26RFa expression (Seifinejad *et al*, 2023), these findings support the view that 26RFa signaling preferentially regulates orexin expression. The high proportion of 26RFa/orexin neurons co-expressing GPR103 is compatible with a paracrine or autocrine mechanism within this neuronal population.

Based on our findings and previous evidence that orexin neurons are activated by hypoglycemia and inhibited by glucose, we propose a working model in which the metabolic consequences of activating LHA 26RFa/orexin neurons depend on the prevailing glycemic state. Within this framework, coordinated transcriptional regulation of the 26RFa and orexin peptidergic systems may contribute to shaping the physiological response of these neurons during metabolic challenges, thereby providing a potential mechanism linking neuronal activity to systemic glucose homeostasis.

In summary, our study identifies a previously unrecognized population of LHA neurons co-expressing 26RFa and orexin that contributes to the regulation of systemic glucose homeostasis. Our findings demonstrate coordinated interactions between these two peptidergic systems at both the functional and transcriptional levels, reveal distinct contributions of 26RFa and orexin signaling to the metabolic response elicited by neuronal activation, and support a role for 26RFa signaling in the regulation of orexin expression. Together, these results provide new insight into the functional organization of hypothalamic peptidergic circuits involved in glucose homeostasis and establish a framework for future studies aimed at defining the mechanisms by which neuronal activity, glycemic state and neuropeptidergic signaling interact to regulate systemic glucose metabolism.

## Data and resource availability

### 1 Data sharing

All data generated or analyzed during this study are included in the published article (and its online supplementary files).

### 2 Resource sharing

No applicable resources were generated or analyzed during the current study.

## Author Contributions

M.D., N.C., and M.P. contributed to the study design and interpretation, and wrote the manuscript. M.D., C.L., S.T., Y.C., A.A., performed the in vivo experiments on mice. M.D., and C.L. contributed to the PCR experiments. M.D. and S.T. performed and analysed the RNAscope experiments. M.D., C.L., M.R., D.G. performed immunohistochimestry experiments. A.B. and E.N. performed the INS assays. B.L. and J.L. produced synthetic 26RFa. J.L. and G.P. revised and approved the final version of the manuscript. N.C. and M.P. are the guarantors of this work, as such, had full access to all the data in the study, and take responsibility for the integrity of the data and the accuracy of the data analysis.

## AI disclosure statement

Generative AI tools were used to assist with language editing and manuscript phrasing. The authors critically reviewed and revised all AI-assisted content and take full responsibility for the final manuscript. AI tools were not used for data generation, data analysis, or interpretation of the experimental results.

## Statement of Ethics

Animal manipulations were performed according to the European Community Council Directive of November 24, 1986 (86:609: EEC), and were conducted by authorized investigators. Our Institutional Animal Use and Care Committee (CENOMAX, Agreement of the Ministry of Research No. 54) approved the experimental procedures (Agreement No.: #38723-2022090217265485).

## Legend to figures

**Supplemental figure 1.**
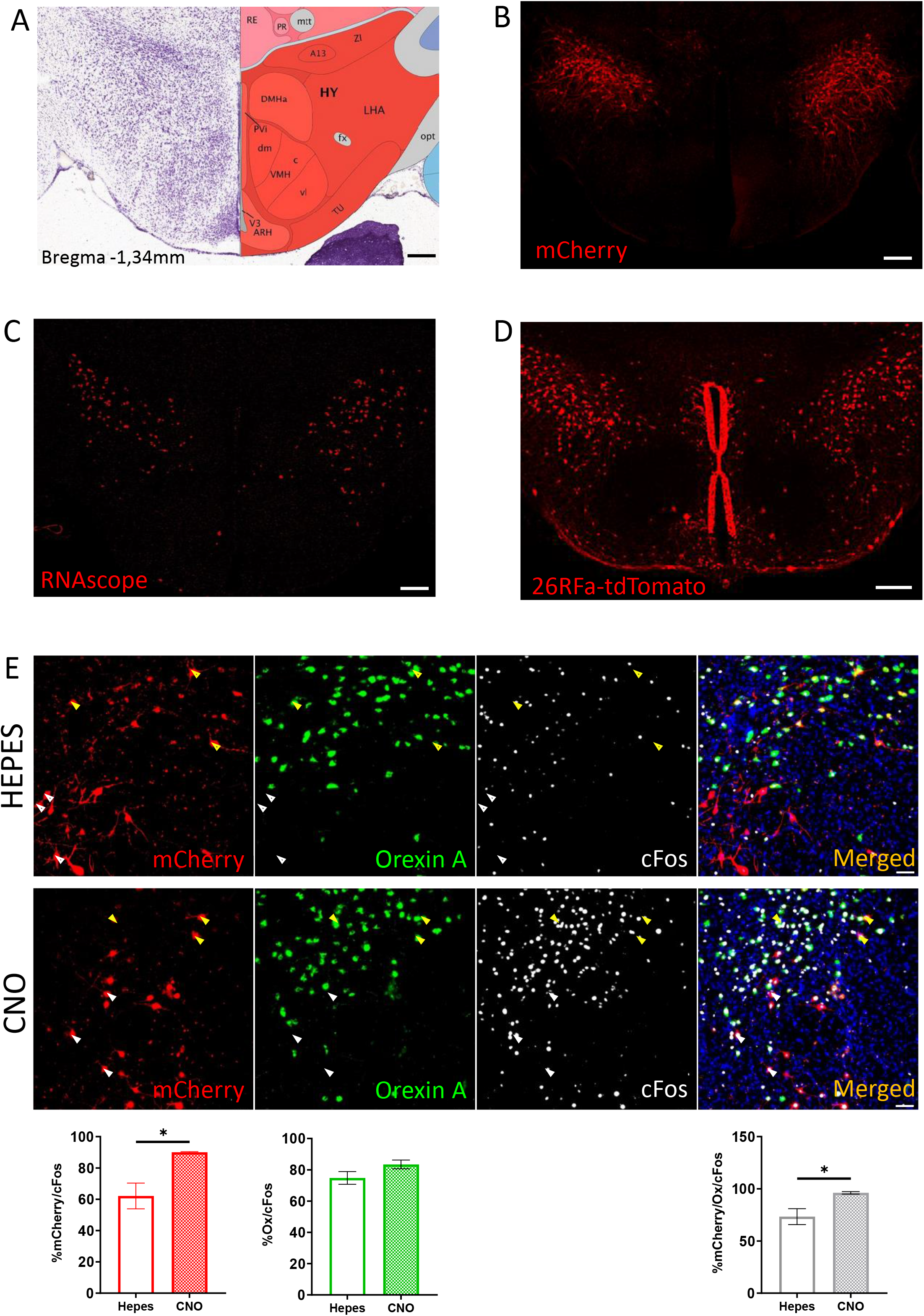
Control experiments showing the localization and the activation of the 26RFa-expressing cells targeted by the chemogenetic approach. **A)** Illustration of Allen Brain Atlas at Bregma −1.34 mm depicting the various nuclei of the hypothalamus. RE : Reuniens Nucleus, PE : Perireuniens, mtt : Mammillothalamic tract, A13 : Dopaminergic A13 group, ZI : Zona incerta, DMHa : Anterior part of dorsomedial hypothalamus, LHA : Lateral hypothalamic area, fx : columns of the fornix, opt : Optic tract, PVi : Intermediate part of periventricular hypothalamus, PVH dm : Dorsomedial part of paraventricular hypothalamic nucleus, VMHc : Central part of ventromedial hypothalamus, VMHvl : Ventrolateral part of ventromedial hypothalamus, TU : Tuberal nucleus, ARH : Arcuate hypothalamic nucleus, V3 : third ventricule. **B)** Panoramic view of a hypothalamic section showing that the expression of the modified muscarinic receptor, vizualized through mCherry fluorescence following stereotaxic AAV injection, is confined to the LHA. **C)** Panoramic view of a hypothalamic section showing that the distribution of prepro-26RFa (red spots) mRNA in the LHA, using the RNAscope *in situ* hybridization approach, overlaps with that of the AAV expression observed in B. **D)** Panoramic view of a hypothalamic section of a 26RFa-tdTomato mouse showing that the main population of 26RFa neurons localized in the LHA overlaps with the mapping depicted in A and B. **E)** Panoramic views of LHA sections showing c-Fos immunoreactivity (grey), orexin-A immunoreactivity (green), and mCherry labeling (red) in the hypothalamus of mice following chemogenetic activation of 26RFa neurons with CNO, compared to basal activity under HEPES (control) treatment. Bar charts show the effect of CNO administration on the percentage of activated neurons among those targeted by the chemogenetic approach (red), the percentage of activated orexin-A-expressing neurons (green), and the percentage of activated neurons co-expressing orexin-A and targeted by the chemogenetic approach (yellow). Data represent the mean ± SEM from 3 mice per condition, with 8–9 hypothalamic sections analyzed per animal. Scale bars: 200µm. \**p*<0.05.

**Supplemental figure 2.**
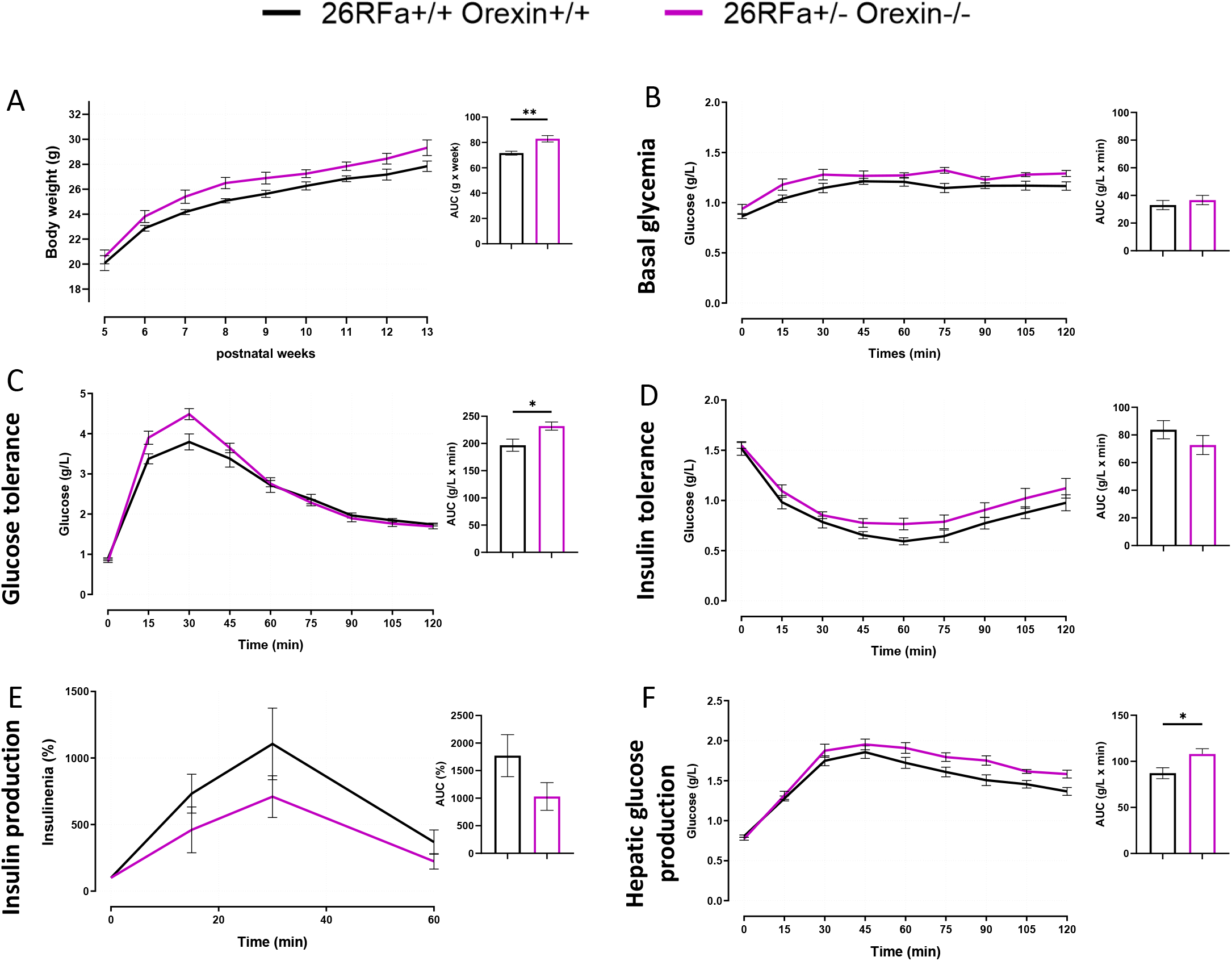
Glycemic phenotype of 26RFa^+/-^;Orexin^-/-^mice. **A)** Body weight evolution from postnatal week 5 to 13 in 26RFa^+/-^;Orexin^-/-^ mice (pink curve) compared to wild-type controls (black curve). **B, C, D, F)** Plasma glucose levels under basal conditions (**B**), during an intraperitoneal (i.p.) glucose tolerance test (**C**), an insulin tolerance test (**D**) and an i.p. pyruvate tolerance test (**F**) in 26RFa^+/-^;Orexin^-/-^ mice (pink curves). **E)** Pasma insulin levels during an i.p. glucose tolerance test in 26RFa^+/-^;Orexin^-/-^ mice (pink curve). \**p*<0.05; \*\**p*<0.01.

**Supplemental figure 3.**
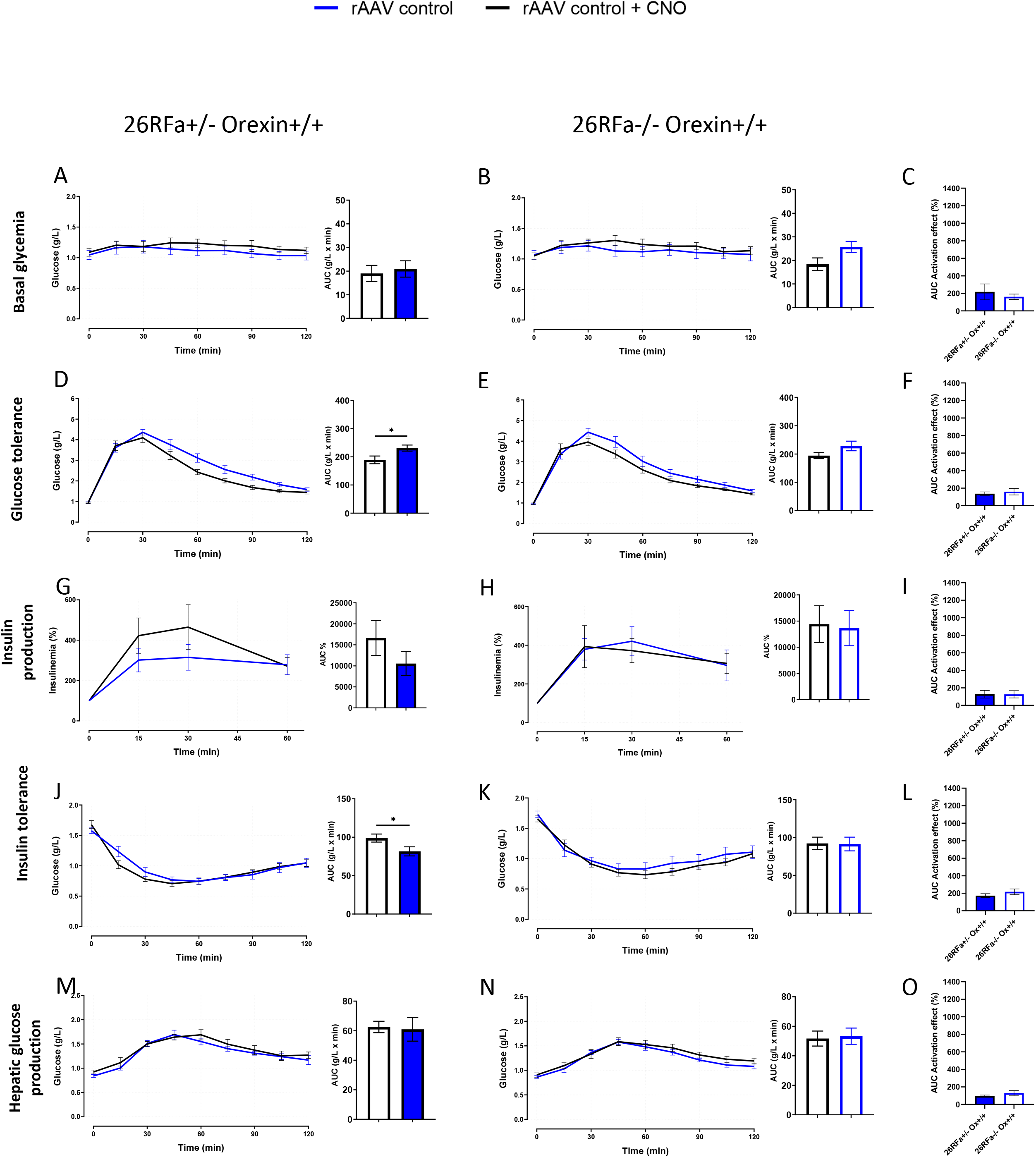
Control experiments showing the effect of the chemogenetic ligand clozapine N-oxide on glucose homeostasis in 26RFa+/- and 26RFa-/- mice injected with the empty AAV. Effects of i.p. administration of clozapine N-oxide (CNO; blue curves) compared to the injection of vehicle (HEPES, black curves) on glucose homeostasis in 26RFa^+/-^ mice **(A, D, G, J, M)** or in 26RFa^-/-^ mice **(B, E, H, K, N)** expressing the control empty AAV in 26RFa neurons. **A, B)** Effect of CNO injection on plasma glucose levels in the basal condition showing the lack of effect of the chemogenetic ligand. **D, E)** Effect of CNO injection on plasma glucose levels during an i.p. glucose tolerance test showing a modest hyperglycemic effect of the chemogenetic ligand. **G, H)** Effect of CNO injection on plasma insulin levels during an i.p. glucose tolerance test showing the lack of effect of the chemogenetic ligand. **J, K)** Effect of CNO injection on glucose levels during an insulin tolerance test showing the lack of effect of the chemogenetic ligand. **M, N)** Effect of CNO injection on glucose levels during an i.p. pyruvate tolerance test showing the lack of effect of the chemogenetic ligand. **C, F, I, L, O)** Area under the curve (AUC) representing the effect of CNO (%). To isolate the specific effect of CNO activation, glycemia values at each time point were normalized to the corresponding values from the non-activated condition. This normalization yielded a relative activation curve (% of control), from which the AUC was calculated to represent the overall activation effect of CNO. Graphs and AUC represent mean ± SEM of 9 mice with each mouse being its own control. \**p*<0.05.

